# Nutritional Regulation of DN1a-Dh44 Signaling Modulates Sleep Across the Lifespan

**DOI:** 10.64898/2026.08.19.745756

**Authors:** Kaitlyn Acklin, Prabriti Neupane, Nabamita Halder, Matthew Li, Amy R. Poe

## Abstract

Across species, sleep amount and timing are tightly linked to the nutritional environment. While early life sleep and sleep in mature organisms are both dramatically influenced by reductions in the dietary environment, the mechanisms linking nutritional cues to conserved sleep-regulatory circuitry are not well understood. Using both early 3^rd^ instar (L3) *Drosophila* larvae and adults, we examined the plasticity of sleep responses under shifting nutrient environments across the lifespan. We find that L3 larvae and adults exhibit changes in sleep duration in low sugar environments with L3 showing a loss of sleep-wake rhythms that can be rescued with additional nutrients. We show that larval and adult sleep plasticity is regulated by *CCHamide-1* signaling between DN1a and Dh44 neurons and glucose metabolic genes in Dh44 neurons. Additionally, our data indicate that sleep plasticity is not dependent on anatomical and functional connectivity between clock-arousal circuitry, suggesting that peptidergic signaling alone is sufficient for diet-dependent sleep regulation. Finally, we demonstrate that Dh44 neurons in both L3 larvae and adults adjust mRNA levels of *CCHamide-1 receptor* (*CCHa1-R*) in response to changes in dietary sugar. Together, our findings suggest that organisms utilize conserved molecular signaling pathways across the lifespan to dynamically regulate their sleep in a changing environment.

## Introduction

Organisms demonstrate a remarkable ability to adjust their sleep in response to a continuously changing environment^1,2^. These changes in sleep timing are likely driven by ecological pressures as time spent sleeping limits the ability to forage, mate, or avoid predation. However, sleep is essential for many physiological processes including metabolic signaling, neural development, and long-term memory formation^3,4^. Therefore, precise regulation of sleep and alignment with the environment is necessary for maintaining overall metabolic homeostasis.

Many species including *Drosophila* modulate their sleep in response to food availability^5–12^. For example, many animals increase sleep after feeding^1^. Conversely, under starvation conditions, most organisms severely limit their sleep to increase the chances of acquiring food^5,7,11^. *Drosophila melanogaster* reduce total sleep duration and increase their activity in periods of starvation^7^. This response is regulated by multiple neurons and pathways including clock neurons, adipokinetic hormone (AKH), octopaminergic neurons, *Diuretic hormone* (Dh44) neurons, and insulin signaling pathways^7,9,13–17^. Finally, reductions in dietary sugar content alter sleep architecture with flies re-partitioning their sleep, increasing the number of sleep bouts *without* altering total sleep, activity, or sleep latency^5,6^ in low sucrose environments. Altogether, these studies indicate that sleep is precisely regulated in response to rapidly changing nutritional environments.

However, these prior studies examining the interactions between alterations in the nutritional environment and sleep have focused exclusively on mature organisms. Yet, sleep plays an important role in brain development as sleep and circadian disruptions early in life negatively impact adult behavior, neuronal morphology, and overall physiology^18–21^. Indeed, in developing organisms, rapid adjustments in sleep timing and foraging in response to nutrient restriction are critical as both neurodevelopment and whole-body tissue development are metabolically taxing processes. In *Drosophila* larvae, behavioral rhythm development is directly influenced by the organism’s nutritional and metabolic status. Sleep-wake and feeding rhythms emerge at the early 3^rd^ instar (L3) stage in *Drosophila*^22,23^. L3 larvae exhibit diurnal sleep and feeding patterns similar to adult flies (increased sleep and decreased feeding at night); the L3 sleep and feeding patterns require a molecular clock, as they are absent in circadian clock mutants^22^. Sleep-wake rhythm development is governed by the formation of a circuit between DN1a clock neurons and Dh44 neurons in the *pars intercerebralis* (PI)^22^. Finally, sleep-wake rhythms are regulated by release of the neuropeptide *CCHamide-1* (*CCHa1*) from DN1as, activating Dh44 arousal-promoting neurons through the *CCHamide-1 receptor* (*CCHa1-R*)^22^. While L3 raised on regular sugar diets (8% glucose) exhibit diurnal differences in sleep, L3 raised on low sugar (1.2% glucose) develop normally, but do not develop sleep-wake rhythms or form the DN1a-Dh44 circuit^22,23^. While adult DN1a and Dh44 neurons regulate sleep in response to environmental disruptions such as starvation^9^ and cold temperatures^24^, it is not known if these neurons act through similar peptidergic signaling pathways across the lifespan to regulate sleep under changing nutritional conditions. Therefore, examining sleep in both larvae and adults provides an opportunity to examine how nutritional signals influence clock-arousal circuit functioning and cellular signaling across the lifespan.

Here, we identify conserved molecular mechanisms underlying nutrient-dependent adaptations in sleep in both larvae and adults. We demonstrate that clock-arousal signaling remains plastic across the lifespan as a mechanism to tune the strength of circadian control over sleep timing. Both L3 and adult flies adapt their sleep in response to changes in the nutritional environment. Additionally, this adaptive response to nutrient changes is regulated by *CCHamide-1* signaling between DN1a clock neurons and Dh44 neurons in both L3 and adult flies. We further demonstrate that both larval and adult Dh44 neurons utilize glucose metabolic genes to regulate diet-induced sleep responses. Finally, we find that larval and adult diet-induced sleep responses are not due to the re-establishment of DN1a-Dh44 circuit connections but are likely due to molecular changes in CCHa1-R mRNA localization in the Dh44 neurons themselves. Together, our data suggests that both DN1a-Dh44 signaling and changes in CCHa1-R subcellular localization allow Dh44 neurons to become competent to receive clock-driven cues and ultimately regulate sleep in response to the levels of dietary sugar in the environment.

## Results

### Sleep regulation is nutritionally plastic across the lifespan

The emergence of sleep-wake rhythms in early 3^rd^ instar (L3) *Drosophila* larvae is regulated by the organism’s energetic and nutritional status^23^. Larvae developing on low glucose (L.G.) (1.2% glucose) diets or forced to adopt immature feeding strategies undergo normal developmental rates, but do not exhibit diurnal changes in sleep^23^. To determine if the loss of sleep-wake rhythms in low sugar diets is permanent, we examined whether re-supplying nutritional sugar to developing larvae restores diurnal rhythms in sleep. We found that moving early L3 raised on L.G. to regular sugar (8% glucose) diets for a period of 4 hours is sufficient to restore sleep-wake rhythms with larvae showing increases in total sleep duration, sleep bout number, and bout length in the subjective evening (Circadian Time [CT] 13) compared to the subjective morning (CT1) (Figure 1A and Figures S1A-B)). However, shorter periods of time (1 or 2 hours) on regular diets do not restore sleep-wake rhythms (Figures 1B-C and Figures S1C-F). Critically, moving early L3 raised on L.G. to L.G. plates (handling controls) for the different time periods failed to restore diurnal differences in sleep duration, sleep bout number, and bout length (Figures 1A-C and Figures S1A-F). Together, these data suggest that developing larvae can modulate sleep duration at specific times of day in response to changes in the nutritional environment.

**Figure 1:**
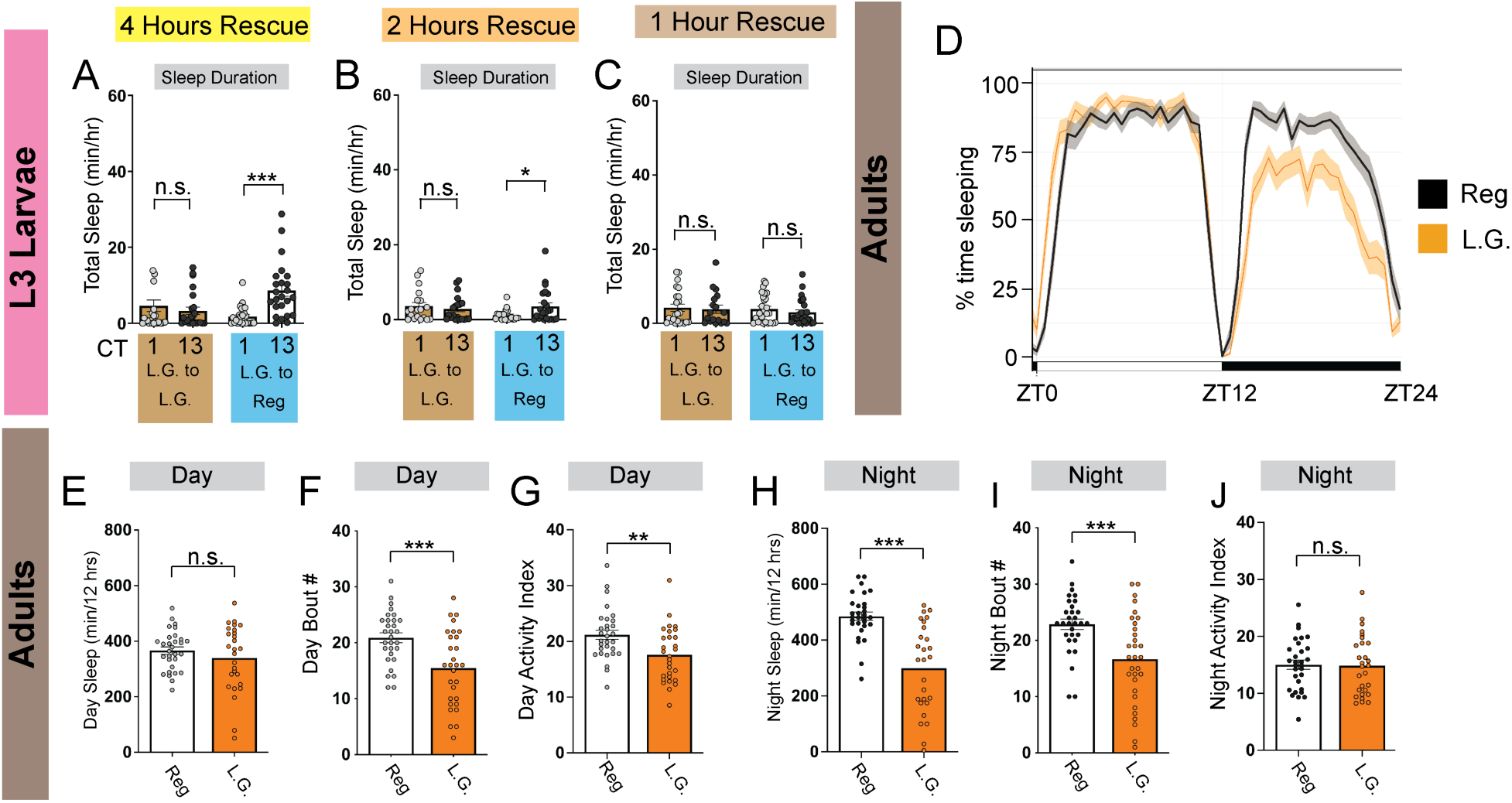
Sleep is nutritionally plastic across the lifespan. (A-C) Sleep duration at CT1 and CT13 in L3 raised on low glucose (L.G.) and moved to regular glucose diets (L.G. to Reg) or moved to L.G. (L.G. to L.G.) for 4 hrs (A), 2 hrs (B), and 1 hr (C). (D-J) Sleep measures for adult flies raised on regular glucose and placed on either low glucose (L.G.) or regular glucose (Reg) for 3 days. (D) Sleep trace depicts sleep amount (%) in rolling 30 min bins across day (ZT0-12, light) and night (ZT12-24, dark). Sleep trace in iso^31^ flies on either L.G. or Reg diets. (E-G) Average daytime sleep duration (E), average number of daytime sleep bouts (F), daytime activity index (G), average nighttime sleep duration (H), average number of nighttime sleep bouts (I), and nighttime activity index (J). Activity index represents the number of infrared beam breaks per minute of waking activity. Sleep experiments were run with multi-beam DAM monitors. A-C, n=18-25 larvae; D-J, n=28-30 flies. Two-way ANOVAs followed by Sidak’s multiple comparison test (A-C); unpaired two-tailed Student’s *t*-test (E-J). For this and all other figures unless otherwise specified, data are presented as mean ± SEM; n.s., not significant; \**P* < 0.05, \*\**P* < 0.01, and \*\*\**P* < 0.001.

A low nutrition environment is also associated with alterations to sleep organization throughout the 24-hour day in adult flies^5^, similar to what we observe in larvae. We next asked whether acute exposure to our low sugar paradigm caused changes in sleep in mature flies. We raised flies on regular glucose diets and then placed newly eclosed flies on either L.G. or regular diets for 5 days. We then examined sleep in L.G. or regular glucose *Drosophila* Activity Monitoring (DAM) tubes using high resolution multi-beam monitors for 3 days. We found that flies on L.G. showed dramatic reductions in sleep duration during the night compared to flies on regular diets (Figures 1D-J and Figures S1G-H). These reductions in nighttime sleep are driven by decreases in sleep bout number, suggestive of more fragmented sleep (Figures 1E-J). While flies on L.G. show reductions in daytime sleep bout number and locomotor activity (Figures 1F-G), daytime sleep duration was not affected (Figure 1E). Additionally, flies on L.G. do not show differences in nighttime locomotor activity during periods of waking, suggesting that overall impairments of motor functioning are not driving nighttime sleep loss (Figure 1J). Altogether, these findings indicate that organisms across the lifespan adapt their sleep in response to reductions in the nutritional environment.

### CCHamide-1 Signaling is necessary for sleep plasticity

The emergence of larval sleep wake rhythms in L3 relies on release of *CCHamide-1* (*CCHa1*) from DN1as to *CCHa1mide-1 receptor* (*CCHa1-R*) on Dh44 neurons^22^. Interestingly, raising larvae on L.G. disrupts the competency of Dh44 neurons to receive *CCHamide-1* signals^23^, suggesting that the nutritional environment influences DN1a-Dh44 communication. To next determine whether *CCHamide-1* signaling is necessary for the restoration of sleep-wake rhythms in our shifting nutrient paradigms, we knocked down *CCHa1* in larval DN1as using *cry-Gal4 pdfGal80* and examined sleep rhythms in L3 raised on L.G. and moved to regular diets for 4 hours. We found that *CCHa1* knockdown disrupted the restoration of sleep-wake rhythms in L3 on regular diets. Instead, *CCHa1* knockdown larvae showed no difference in sleep duration, sleep bout number, or bout length between CT1 and CT13 after being on regular sugar diets for 4 hours (Figure 2A and Figures S2A-B). However, genetic controls showed the expected increase in sleep duration, sleep bout number, and bout length at CT13 after 4 hours on regular diets (Figure 2A and Figures S2A-B). We next performed the converse experiment and knocked down *CCHa1-R* in larval Dh44 neurons and examined sleep rhythms in L3 raised on L.G. and moved to regular diets for 4 hours. As expected, we observed no difference in sleep duration, bout number, or bout length between CT1 and CT13 in the RNAi-expressing larvae (Figure 2B and Figures S2C-D), indicating that *CCHamide-1* signaling between DN1a and Dh44 neurons is necessary for the rescue of larval sleep rhythms under shifting nutrient conditions.

**Figure 2:**
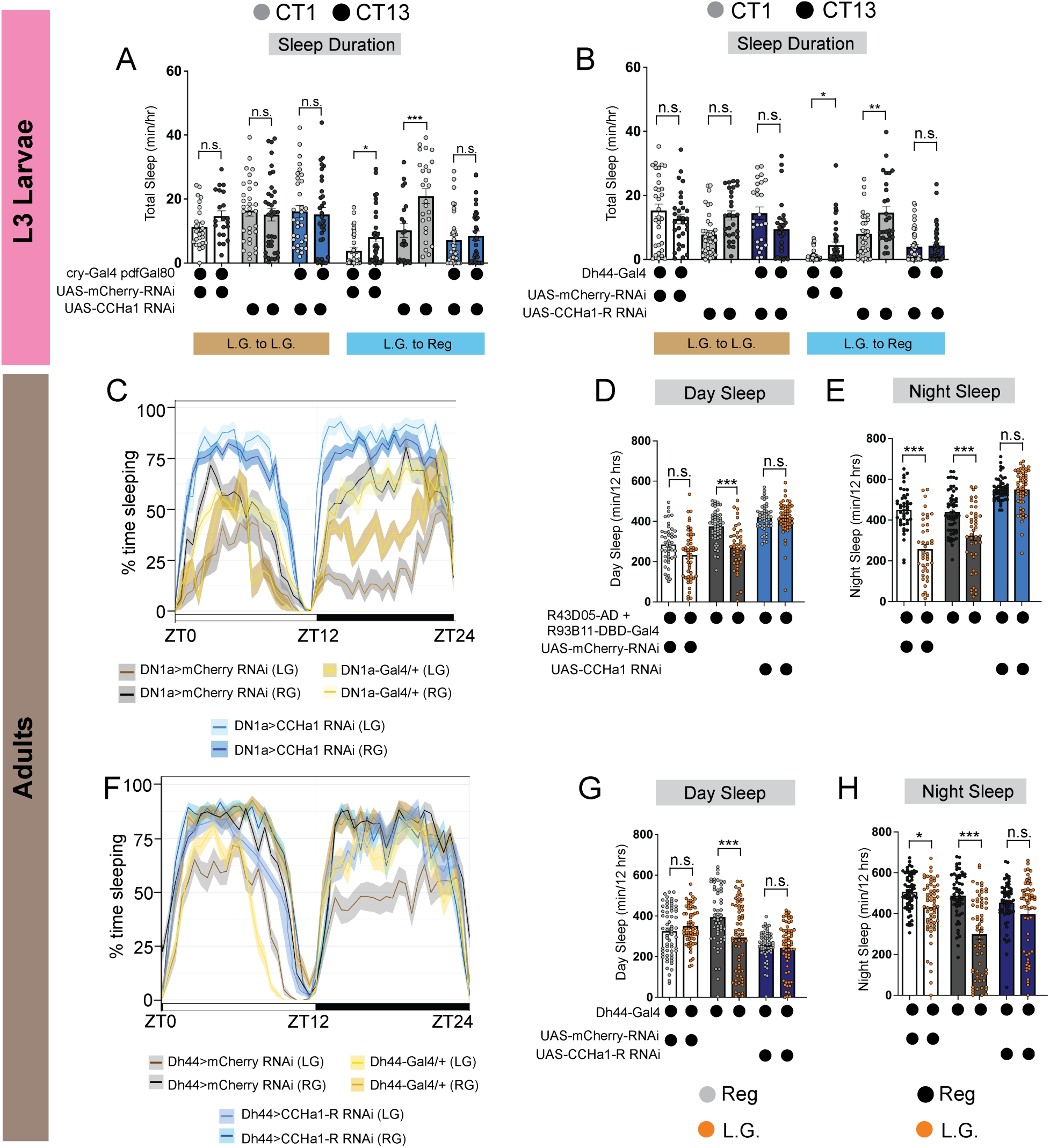
CCHamide-1 signaling is necessary for sleep plasticity. (A-B) Sleep duration in L3 expressing UAS-CCHa1-RNAi with cry-Gal4 pdfGal80 (A) and UAS-CCHa1-R-RNAi with Dh44-Gal4 (B) and genetic controls at CT1 and CT13 in diet shifting paradigms. Larvae were on either L.G. or Reg diets for 4 hours. (C) Sleep trace in adults expressing UAS-CCHa1-RNAi with DN1a-specific Gal4 and genetic controls on low glucose (L.G.) or regular glucose (Reg) diets. (D-E) Average daytime (D) and nighttime (E) sleep in adults expressing UAS-CCHa1-RNAi with DN1a-specific Gal4 and genetic controls. (F) Sleep trace in adults expressing UAS-CCHa1-R-RNAi with Dh44-Gal4 and genetic controls on L.G. or Reg diets. (G-H) Average daytime (G) and nighttime (H) sleep in adults expressing UAS-CCHa1-R-RNAi with Dh44-Gal4 and genetic controls. Orange dots indicate flies on L.G. diets. Adult sleep experiments were run with multi-beam DAM monitors. A-B, n=25-47 larvae; C-E, n=44-54 flies; F-H, n=55-64 flies. Two-way ANOVAs followed by Sidak’s multiple comparison test (A-B); One-way ANOVAs followed by Tukey’s multiple comparisons test (D-E and G-H).

We next asked if *CCHamide-1* signaling between DN1a and Dh44 neurons is necessary for the reduction in sleep seen in adults on L.G. diets. Using a DN1a-specific split Gal4 driver (R43D05-AD + R93B11-DBD-Gal4)^25^, we knocked down *CCHa1* in adult DN1as and examined sleep in flies raised on regular glucose diets and then placed on either L.G. or regular diets for 5 days. We found that *CCHa1* knockdown flies did not show a reduction in nighttime sleep duration on L.G. diets compared to knockdown flies on regular glucose diets (Figures 2C-E). Additionally, *CCHa1* knockdown flies did not show any differences in daytime sleep duration, sleep bout number, or average sleep bout length on L.G. diets compared to knockdown flies on regular diets (Figures 2C-E and Figures S3A-F). Genetic controls showed the expected reduction in nighttime sleep duration and nighttime sleep bout number on L.G. diets (Figures 2C-E and Figures S3A-F). Finally, expressing *CCHa1-R* in adult Dh44 neurons also eliminated the L.G.-induced reductions in sleep duration and sleep bout number during the night compared to both flies on regular diets and genetic controls on L.G. diets (Figures 2F-H and Figures S4A-F). Combined with our larval manipulations (Figures 2A-B and Figures S2A-D), these data indicate that *CCHamide-1* signaling between DN1a and Dh44 neurons plays a conserved role in regulating diet-induced sleep responses.

### Dh44 neurons require glucose metabolic genes for sleep plasticity

Finally, our prior work showed that Dh44 neurons drive sleep-wake rhythm development on regular diets through the actions of glucose metabolic genes^23^. Our data suggest that similar to their adult counterparts^9,16,26,27^, larval Dh44 neurons may sense the nutritional environment. To test if this is the case in L3 raised on L.G. and moved to regular diets for 4 hours, sleep duration at CT1 and CT13 was assessed with knockdown of glucose metabolic genes (*Hexokinase-C*, *Glucose transporter 1,* and *Pyruvate kinase*) in Dh44 neurons. We found that knockdown of the glucose metabolism genes, *Glut1*, *Hex-C* and *PyK*, in Dh44 neurons prevented the emergence of diurnal rhythms in sleep duration and bout number when L3 were shifted from L.G. to regular diets (Figures 3A-C and Figures S5A-F). However, genetic controls showed the expected increases in sleep duration, sleep bout number, and bout length at CT13 after 4 hours on regular diets (Figures 3A-C and Figures S5A-F). These findings suggest that larval Dh44 neurons require components of the glycolysis pathway for rapid changes in sleep consolidation in response to changing glucose levels.

**Figure 3:**
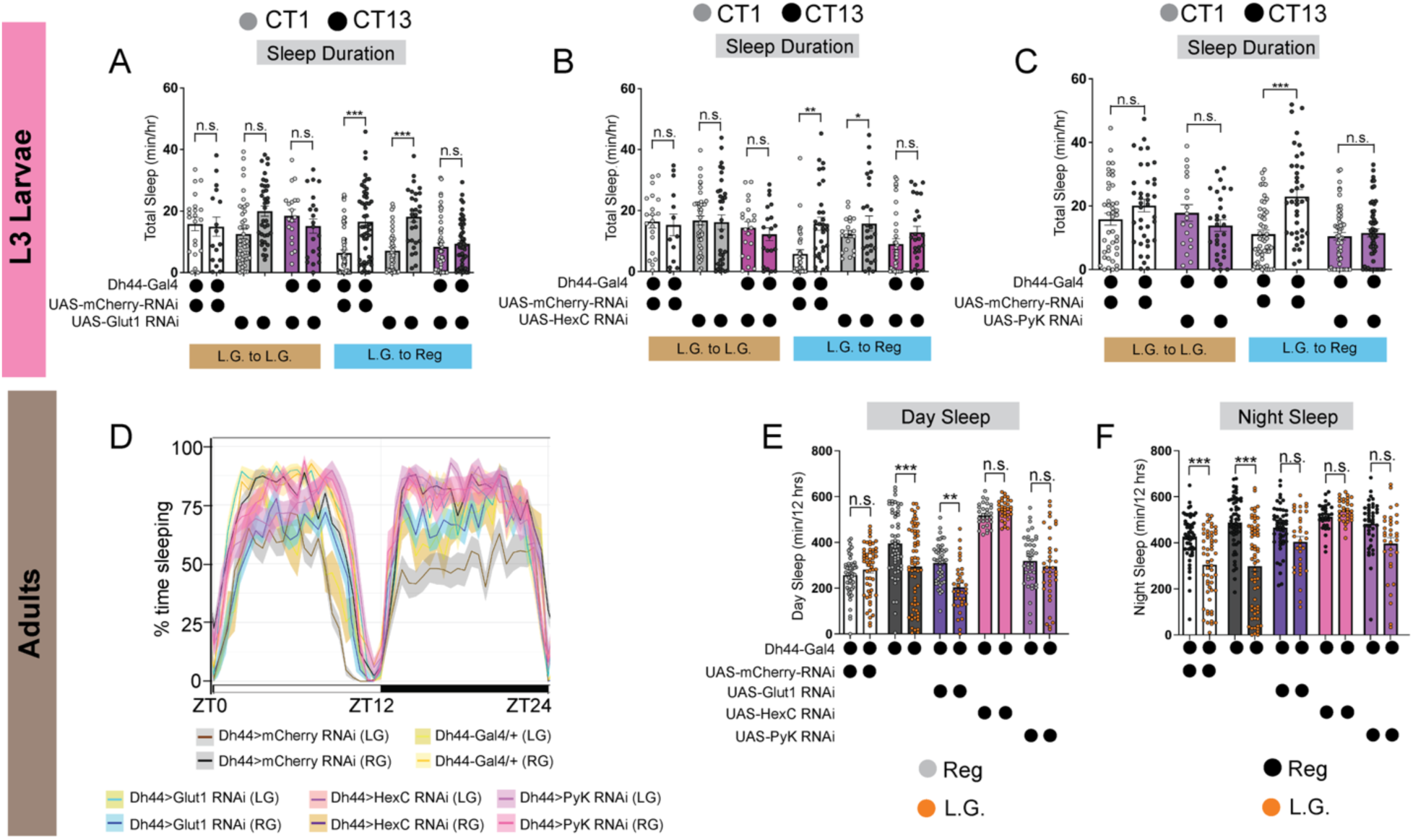
Dh44 neurons require glucose metabolic genes for sleep plasticity. (A-C) Sleep duration in L3 expressing UAS-Glut1-RNAi (A), UAS-Hex-C-RNAi (B), and UAS-Pyk-RNAi (C) with Dh44-Gal4 and genetic controls at CT1 and CT13 in diet shifting paradigms. Larvae were on either L.G. or Reg diets for 4 hours. (D) Sleep trace in adults expressing UAS-Glut1 RNAi, UAS-HexC RNAi, and UAS-PyK RNAi with Dh44-Gal4 and genetic controls on L.G. or Reg diets. (E-F) Average daytime sleep duration (E) and average nighttime sleep duration (F) in adults expressing UAS-Glut1 RNAi, UAS-HexC RNAi, and UAS-PyK RNAi with Dh44-Gal4 and genetic controls. Orange dots indicate flies on L.G. diets. Adult sleep experiments were run with multi-beam DAM monitors. A-C, n=25-54 larvae; D-F, n=30-60 flies. Two-way ANOVAs followed by Sidak’s multiple comparison test (A-C); One-way ANOVAs followed by Tukey’s multiple comparisons test (E-F).

To next determine if adult Dh44 neurons require glucose metabolism genes for the loss of sleep on L.G. diets, we examined sleep in *Glut1*, *HexC*, and *PyK* knockdown flies raised on regular glucose diets and then placed on either L.G. or regular diets for 5 days. We found that knockdown of the glucose metabolism genes in adult Dh44 neurons prevented the expected reduction in nighttime sleep duration and sleep bout number on the L.G. diet compared to both flies on regular diets and genetic controls on L.G. diets (Figures 3D-F and Figures S6A-F). The *Glut1* knockdown flies did show a small, but significant reduction in daytime sleep duration and average daytime sleep bout number on the L.G. diet (Figures 3D-F and Figures S6A-D) possibly due to insufficient knockdown of gene expression. However, sleep bout length and locomotor activity were not affected in the RNAi-expressing flies on the L.G. diets (Figures S6A-F). Together, our data suggest that Dh44 neurons utilize glucose metabolism genes to regulate sleep in response to the diet in both larvae and adult flies.

### DN1a-Dh44 circuit formation is not nutritionally plastic

In L3 larvae, sleep-wake rhythm development depends on anatomical and functional connections forming between DN1a and Dh44 neurons^22,23^. These functional connections are developmentally plastic as larvae raised on L.G. diets do not form circuit connections between the central clock and arousal neurons^23^. We next examined synaptic connections between DN1a and Dh44 neurons in the setting of shifting nutritional conditions using neurexin-based GFP reconstitution across synaptic partners (GRASP)^28–30^. Using *cry*-Gal4 (larval s-LNv/DN1a-specific driver) with *Dh44-*LexA to express independent GRASP components, we observed the expected reconstituted GFP signal around the cell body and dendrites of Dh44 neurons in L3 raised on regular diets (Figures 4A-A’)^22^. However, we did not observe GFP signal in L3 raised on L.G. and moved to L.G. for 4 hours (Figures 4B-B’). Finally, we observed GFP reconstitution with reduced signal intensities around the cell body of Dh44 neurons in L3 raised on L.G. and moved to regular diets for 4 hours (Figures 4C-C’). Indeed, quantification of GFP-positive puncta indicates that fewer synaptic connections form between DN1a and Dh44 neurons in shifting nutritional conditions (Figure 4D), suggesting that anatomical connectivity is only partially restored with the addition of nutritional sugar.

**Figure 4:**
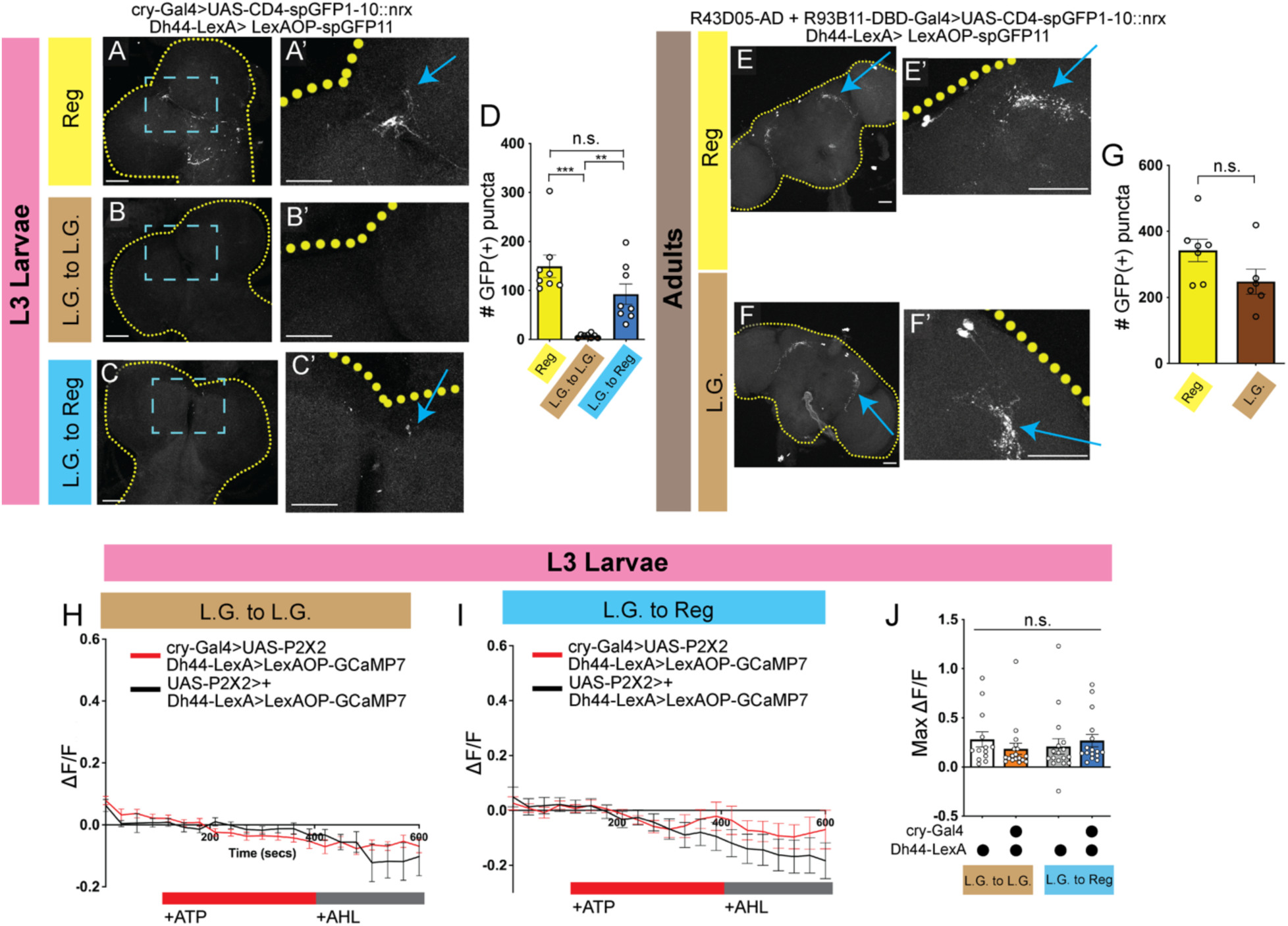
DN1a-Dh44 circuit formation is not nutritionally plastic. (A-C’) Neurexin-based green fluorescent protein (GFP) reconstitution (GRASP; GFP reconstitution across synaptic partners) between DN1as (cry-Gal4) and Dh44 neurons (Dh44-LexA) in larvae raised on regular food (A), raised on low glucose (L.G.) and moved to L.G. (B), or raised on L.G. and moved to regular food (C) after 4 hours. Higher magnification of region of interest in (A’), (B’), and (C’). (D) Quantification of number of GFP-positive puncta in region of interest in A’, B’, and C’. (E-F’) Neurexin-based GRASP between adult DN1as (R43D05-AD + R93B11-DBD) and Dh44 neurons (Dh44-LexA) in adults on either regular glucose or L.G. diets. Higher magnification of region of interest in (E’) and (F’). (G) Quantification of number of GFP-positive puncta in region of interest in E and F. (H-I) GCaMP7 signal in Dh44 neurons with the activation of DN1a neurons in L.G. to L.G. diet shifts (H) or L.G. to Regular diet shifts (I). Larvae were on either L.G. or Reg diets for 4 hours. The red bar indicates adenosine 5’-triphosphate (ATP) application and the gray bar indicates artificial hemolymph (AHL) application. (J) Maximum GCaMP change (ΔF/F) for individual cells in L.G. to L.G. (H) or L.G. to Reg (I). For A-F’, yellow dotted lines = brain; blue arrows = GRASP signal. A-C’, n=8-10 brains; E-F’, n=8 brains; H-J, n=16-18 cells, 10 brains. One-way ANOVAs followed by Tukey’s multiple comparisons test (D and J); unpaired two-tailed Student’s *t*-test (G). Scale bars, 25 µm for A-C’ and 50 µm for E-F’.

Next, to determine if adult DN1a-Dh44 connectivity is also influenced by the nutritional environment, we raised flies on regular diets and examined synaptic connections between DN1a and Dh44 neurons using neurexin-based GRASP^28–30^ after 5 days on either regular or L.G. diets. Using a DN1a-specific split Gal4 driver (R43D05-AD + R93B11-DBD-Gal4)^25^ with *Dh44-*LexA to express independent GRASP components, we observed GFP reconstitution around the cell body and dendrites of Dh44 neurons in flies on both regular and L.G. diets (Figures 4E-F’) compared to negative controls (Figure S7A-B’). In line with this, quantification of GFP-positive puncta indicates that flies exhibit similar numbers of synaptic connections between DN1a and Dh44 neurons on both diets (Figure 4G) suggesting that the number of synapses is not affected by loss of nutrients. This surprising result indicates that changes in anatomical connectivity between DN1a and Dh44 neurons do not underlie the observed sleep loss in L.G. diets. Instead, combined with our RNAi knockdown results, our findings suggest that changes in peptidergic signaling alone are sufficient to modulate sleep in response to shifting diets.

Finally, we examined functional connectivity between larval DN1a and Dh44 neurons in the context of shifting nutritional conditions. We expressed ATP-gated P2X2 receptors^31^ in DN1a neurons and GCaMP7 in Dh44 neurons. Our previous work showed that activation of DN1as in L3 raised on regular food resulted in a calcium response in Dh44 neurons^22,23^. As expected, activation of DN1as in L3 raised on L.G. food and moved to L.G. for 4 hours did not elicit a calcium response in Dh44 neurons (Figures 4H-J)^23^. Surprisingly, in L3 raised on L.G. food and moved to regular food for 4 hours, activation of DN1as also did not elicit a calcium response in Dh44 neurons (Figures 4I-J). Thus, while robust sleep-wake rhythms emerge after 4 hours on regular food, their emergence likely relies on non-synaptic communication between DN1a clock and arousal loci.

### CCHa1-R expression in Dh44 neurons is nutritionally plastic

Thus far, our data indicate that both L3 larvae and adult flies adjust their sleep in response to changes in glucose levels in the dietary environment. Additionally, our RNAi knockdown experiments suggest that these adaptations to the levels of glucose depend on *CCHamide-1* signaling between DN1a and Dh44 neurons. However, surprisingly, these changes in sleep are likely not due to changes in anatomical and functional connectivity between DN1a and Dh44 neurons on shifting diets. Previously, we found that raising larvae on L.G. environments disrupts the competency of Dh44 neurons to receive *CCHamide-1* signals^23^. Therefore, we next asked whether larval Dh44 neurons exhibit changes in *CCHa1-R* expression levels in shifting nutrient environments. We performed RNA fluorescent *in situ* hybridization chain reaction (HCR) staining of *CCHa1-R* in Dh44 neurons in L3 raised on regular food, raised on L.G. and moved to L.G. for 4 hours, or raised on L.G. and moved to regular food for 4 hours. We observed a loss of *CCHa1-R* mRNA expression in Dh44 neurons on L.G. diets compared to control diets (Figures 5A-B’’). Surprisingly, moving L3 from L.G. to regular food for 4 hours restored *CCHa1-R* mRNA expression levels in Dh44 neurons (Figures 5C-C’’). Indeed, quantification of *CCHa1-R* expression in Dh44 cell bodies showed an increase in the L.G. to regular food condition compared to regular food alone or L.G. to L.G. handling controls (Figure 5D).

**Figure 5:**
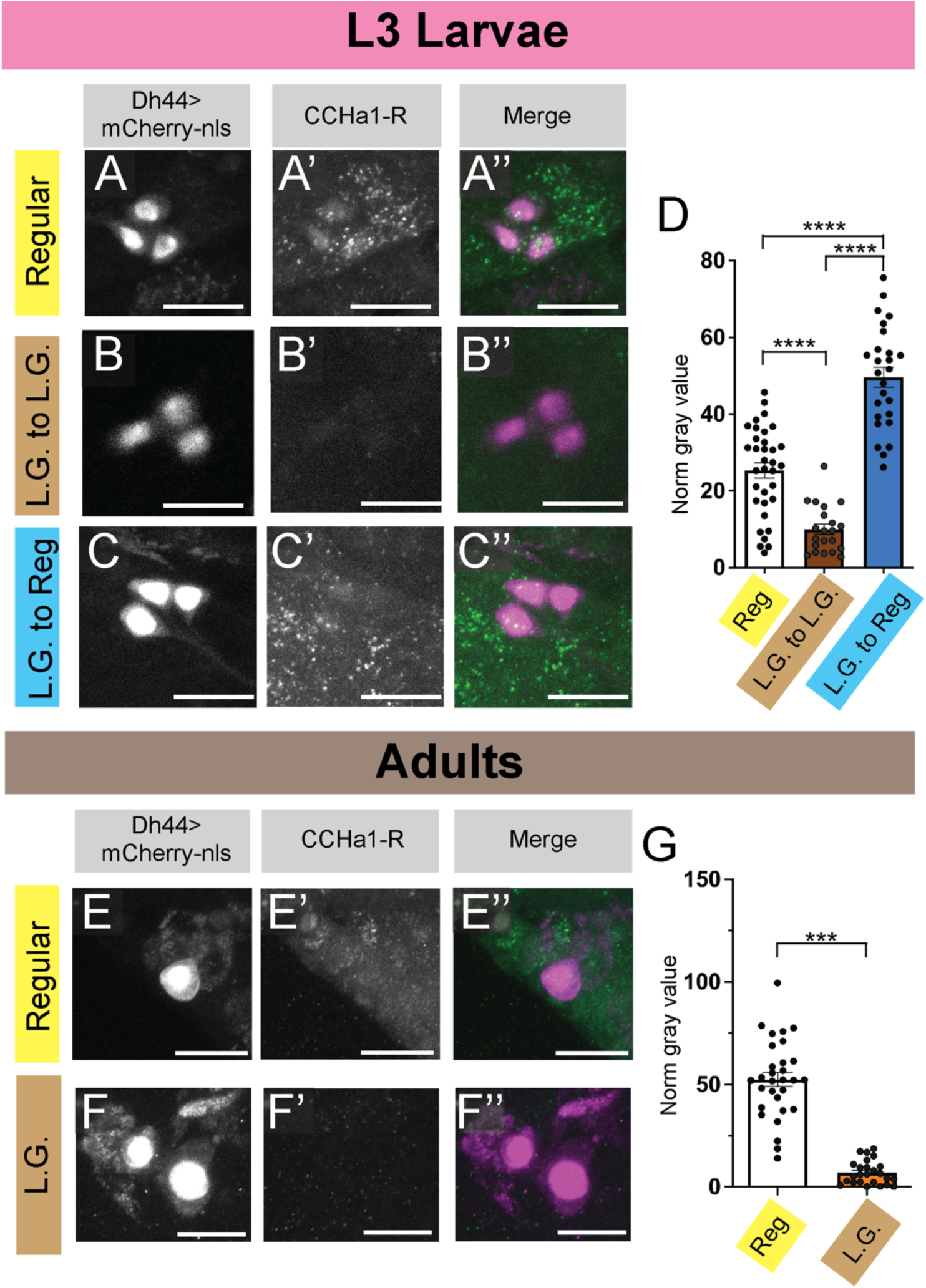
CCHa1-R Expression Changes Under Nutrient Shifting Conditions. (A-C) RNA fluorescent in situ hybridization (HCR) labeling of CCHa1-R in larval Dh44 neurons in larvae raised on regular food (A), raised on low glucose (L.G.) and moved to L.G. (B), or raised on L.G. and moved to regular food (C) after 4 hours. (D) Quantification of HCR staining in larval Dh44 cell bodies in the different diet conditions. (E-F’’) HCR labeling of CCHa1-R in adult Dh44 neurons in flies on either regular food (E-E’’) or low glucose (F-F’’) diets. (G) Quantification of HCR staining in adult Dh44 cell bodies in the different diet conditions. Dh44 neurons are labeled with mCherry-nls and the CCHa1-R HCR probe set is labeled with Alexa-Fluor 488. For D and G, gray value is normalized to background signal. A-D, n=21-34 cells, 8-10 brains; E-G, n=25-30 cells, 10 brains. One-way ANOVA followed by Sidak’s multiple comparison test (D); unpaired two-tailed Student’s *t*-test (G). Scale bars, 25 µm for A-C’’ and 10 µm for E-F’’.

Finally, we examined *CCHa1-R* mRNA expression levels in adults on regular and L.G. food using HCR. We observed a loss of *CCHa1-R* mRNA expression in Dh44 neurons on L.G. diets compared to control diets (Figure 5E-F’’). In support of this, quantification of *CCHa1-R* expression in adult Dh44 cell bodies showed a dramatic reduction compared to the regular diet (Figure 5G), indicating that adult Dh44 neurons modulate *CCHa1-R* expression in response to reductions in dietary sugar. Altogether, our findings indicate that diet-dependent adjustments in sleep are driven by peptidergic signaling between DN1a and Dh44 neurons and changes in *CCHa1-R* expression levels in Dh44 neurons.

## Discussion

The nutritional environment greatly influences sleep in multiple organisms, but the molecular mechanisms linking nutritional cues to sleep-regulatory circuits are not well understood. Here, we demonstrate that adjustments in sleep due to changes in dietary sugar levels depend on communication between clock (DN1a) and arousal (Dh44) neurons across the lifespan. Our data indicate that *CCHamide-1* signaling is necessary for the modulation of sleep in low glucose environments as manipulations of both *CCHa1* in DN1as and *CCHa1-R* in Dh44 neurons disrupt sleep plasticity. Additionally, our data demonstrate that larval and adult Dh44 neurons require glucose metabolic genes to regulate sleep in response to reductions in nutritional sugar levels. Finally, our analysis of DN1a-Dh44 circuitry under shifting nutritional conditions suggest that clock-arousal circuit formation itself is *less* plastic to changing environments. Instead, changes in *CCHa1-R* expression in Dh44 neurons likely underlie diet-dependent sleep responses. These findings suggest that organisms at different life stages regulate sleep in response to changes in nutrient availability through similar, conserved signaling pathways.

Together, with our previously published work^23^, our findings support a model in which changes in neuropeptidergic communication between DN1a and Dh44 neurons drive sleep-wake rhythm development and regulate sleep homeostasis across the lifespan. Our CCHa1-R HCR data suggest that Dh44 neurons promote *CCHa1-R* expression in response to increases in glucose levels in the environment. This increase in *CCHa1-R* expression allows Dh44 neurons to become competent to receive clock-driven cues. This model suggests that rapid changes in sleep in response to shifting nutritional conditions utilize paracrine signaling alone and not circuit-level changes between clock and arousal neurons. In support of this model, our GRASP data suggests that anatomical connections alone are not required for modulating sleep in response to changing nutrients. Finally, our larval P2X2 data indicate that circuit connections between DN1a and Dh44 neurons do not completely form in rapidly changing diets, providing further support for the conclusion that signaling between clock and arousal neurons is a key factor regulating sleep plasticity.

Our findings expand on published studies examining sleep regulation in periods of nutrient scarcity including starvation^7,9^ and low sucrose environments^5,6^. Similar to prior studies^9^, our data validate the role of Dh44 neurons in regulating sleep responses in starvation conditions. However, in contrast to previous studies, our data provide new insights into the possible molecular mechanisms that Dh44 neurons utilize to modulate sleep across the lifespan. While we demonstrate a role for neurons in the brain in regulating sleep, our data does not exclude the possibility that signaling from peripheral tissues or from other populations of clock output neurons are necessary for DN1a-Dh44 communication and sleep regulation in shifting nutrient conditions. *Drosophila insulin-like peptide 2* (*Dilp2)* released from the insulin-producing cells (IPCs) in the PI also modulates starvation-induced changes in adult sleep depth^17^. Interestingly, IPCs express receptors for Dh44 neuropeptides^32^, suggesting that cross-communication between Dh44 neurons and IPCs may be necessary for diet-induced sleep plastic responses. Previous work has also shown that gut-derived *CCHa1* signals regulate adult fly sleep, sleep depth, and feeding^33^ in standard dietary conditions. Therefore, future studies will examine the role of other clock output neuron populations and will determine whether gut-derived signals also modulate sleep responses to reduced nutrient environments.

Finally, our findings imply that changes in the dietary environment have evolutionary conserved roles in regulating sleep. In rodents, chronic low nutrient diets during development disrupt normal sleep-wake patterns and cognitive function in mature offspring^34–36^. As early-life nutrition is also associated with sleep quality, neurodevelopment, and cognitive outcomes in humans, identifying conserved mechanisms that couple nutrient sensing to sleep may provide new insights into how developmental nutritional experiences shape long-term health^37^. However, whether transient changes in the levels of nutrient available during mammalian development have lasting effects on sleep and memory formation across the lifespan is not known. Furthermore, our prior work demonstrated that raising larvae on L.G. diets or forcing larvae to adopt constant feeding strategies (at the expense of sleep-wake rhythms) resulted in a loss of long-term aversive memory (LTM) performance^23^. These studies suggest that adaptations in sleep amount and LTM in response to the nutritional environment might provide immediate benefits to organisms by limiting energetically costly processes. Future work will utilize the larval system to define how memory formation is regulated by the sensing of the changing nutritional environment.

## Acknowledgements

We thank members of the Paré lab for helpful suggestions on the HCR staining. We thank members of the Poe lab and Dan Cavanaugh for helpful discussions and input.

## Author contributions

Conceptualization, K.A., A.R.P.; Investigation, K.A., P.N., N.H., M.L., A.R.P.; Writing – Original Draft, K.A., P.N., and A.R.P.; Writing – Review & Editing, all authors; Project supervision and funding, A.R.P.

## Data and Materials Availability

All data needed to evaluate the conclusions in the paper are present in the paper and/or the Supplementary Materials.

## Materials and Methods

### Fly Stocks

The following lines have been maintained as lab stocks: iso31, Dh44^VT^-Gal4 (VT039046)^38^, cry-Gal4 pdf-Gal80^39^, UAS-GCaMP7f UAS-tdTom^40^, LexAOP-sp-GFP11; UAS-CD4-spGFP1-10::nrxn^28^, and UAS-mCherry RNAi. CCHa1-R RNAi (51168), CCHa1 RNAi (57562), Hex-C RNAi (57404), Glut1 RNAi (40904), PyK RNAi (35218) and UAS-mCherry-nls (38424) were from the Bloomington *Drosophila* Stock Center (BDSC). R43D05-AD, R93B11-DBD-Gal4 was obtained from Dan Cavanaugh.

### Larval rearing and sleep assays

Larval sleep experiments were performed as described previously^22,23,41^. Briefly, molting 3^rd^ instar larvae were placed into individual wells of the LarvaLodge containing 95 µl of 3% agar and 2% sucrose media covered with a thin layer of yeast paste. The LarvaLodge was covered with a transparent acrylic sheet and placed into a Percival dual-chamber incubator at 25°C for imaging. Experiments were performed in the dark.

### LarvaLodge image acquisition and processing

Images were acquired every 6 seconds with an Imaging Source DMK 23UP031 camera (2592 × 1944 pixels, The Imaging Source, USA) equipped with a Fujinon lens (HF12.55A-1, 1:1.4/12.5 mm, Fujifilm Corp., Japan) with a Hoya 49mm R72 Infrared Filter as described previously^22,41^. We used IC Capture 4 (The Imaging Source) to acquire time-lapse images. All experiments were carried out in the dark using infrared LED strips (Ledlightsworld LTD, 850 nm wavelength) positioned below the LarvaLodge.

Images were analyzed using custom-written MATLAB software (see Churgin et al 2019^42^ and Szuperak et al 2018^41^). Temporally adjacent images were subtracted to generate maps of pixel value intensity change. A binary threshold was set such that individual pixel intensity changes that fell below 45 gray-scale units within each well were set equal to zero (“no change”) to eliminate noise. Pixel changes greater than or equal to threshold value were set equal to one (“change”). Activity was then calculated by taking the sum of all pixels changed between images. Sleep was defined as an activity value of zero between frames. For sleep experiments performed at certain circadian times, total sleep in the 2^nd^ hour after the molt to third instar was summed.

### Dietary Manipulations

Fly food was prepared using previously described recipes^23^. For larval experiments, adult flies were placed in an embryo collection cage (Genesee Scientific, cat#: 59-100) and eggs were laid on a petri dish containing Low glucose (1.2% glucose) (L.G.) food. Animals developed on this media for three days. Molting 3^rd^ instar larvae were then selected and moved to either control (8% glucose) or L.G. food for the indicated times.

### Adult Sleep Experiments

Adult flies were raised at 25°C on a 12h:12 h light:dark (LD) cycle on control food. Day 0-1 males were then moved to either control or L.G. vials and group housed for 5 days. Flies were then anesthetized on CO2 pads and loaded into Drosophila Activity Monitoring tubes containing control or L.G. food. Activity was monitored for 3 days using multi-beam DAM monitors. For all experiments, activity was measured in 1 min bins and sleep was defined as 5 minutes of inactivity. Activity indices were measured as number of beam breaks per 1 min of waking activity. Data was analyzed using Rethomics.

### Immunohistochemistry & imaging

Brains were dissected in PBS, fixed in 4% PFA for 20 min at room temperature. Following 3 × 20 min washes in PBST, brains were incubated with primary antibody at 4°C overnight. Following 3 × 20 min washes in PBST, brains were incubated with secondary antibody at 4°C overnight. Following 3 × 20 min washes in PBST, brains were mounted in Vectashield (Vector Laboratories. Primary antibodies included: Rabbit anti-GFP (1:200, A-11122, ThermoFisher Scientific), Mouse anti-GFP (for GRASP dissections) (1:200, EMSCO/Fisher, Cat. No: G6539-.2ML), and Mouse anti-mCherry (for HCR staining) (1:200, Fisher, Cat. No: NC0288633). Secondary antibodies included Alexa Fluor donkey anti-rabbit 488 (1:500, Jackson), goat anti-mouse 488 (1:500, Jackson), and donkey anti-mouse 594 (1:500, Fisher, Cat. No: A32744). Brains were visualized and imaged with a Zeiss LSM 900 confocal microscope. For GRASP quantifications, the number of GFP-positive puncta in each ROI was quantified using the Analyze Particles plug-in in ImageJ. Quantifications were performed blind to experimental condition.

### RNA Fluorescence *in situ* hybridization (RNA-FISH)

Custom probes for CCHa1-R were obtained from Molecular Instruments and labeled with Alexa-Fluor 488. For staining and labeling of Dh44 cell bodies, Dh44-Gal4 was crossed with UAS-mCherry-nls. CCHa1-R was labeled with CCHa1-R probe and 488 hairpins (h1/h2). Staining was performed as described in the Molecular Instruments HCR protocol for whole-mount fruit fly embryos. The entire staining protocol took 4 days. Briefly, larval and adult brains were dissected in Schneider’s media and then fixed in 4.5% paraformaldehyde for 25 minutes. Brains subsequently underwent a series of wash steps in methanol, ethanol, xylene and PBST. Following the wash steps, brains were incubated overnight at 37°C in 0.8 pmol of probe solution. The next day, brains were washed with probe wash buffer and 5x SSCT before incubating overnight at room temperature in the h1/h2 hairpin solution. On day 3, excess hairpins were removed using 5x SSCT washes and brains were incubated overnight at 4°C in primary antibody (mouse anti-mCherry). On day 4, brains were washed with 5x SSCT, incubated for 2 hours at room temperature in secondary antibody (donkey anti-mouse 594), washed again with 5x SSCT, and mounted on slides with Vectashield (Fisher, Cat. No: NC9265087). Brains were visualized and imaged with a Zeiss LSM 900 confocal microscope. *CCHa1-R* expression was quantified in each cell body using the mCherry-nls channel to define the ROI in ImageJ. Then, images were processed using the Process-Smooth command to remove background noise. Finally, the mean gray value in the *CCHa1-R* channel in each ROI was measured. Gray values were normalized to neighboring regions. All analysis was done blind to experimental condition.

### P2X2 imaging

Early 3^rd^ instar larvae were raised on L.G. food and then moved to either control or L.G. food for 4 hours. Live imaging was performed as described previously^22,23^. Briefly, larval brains were then dissected in artificial hemolymph (AHL) buffer consisting of (in mM): 108 NaCl, 5 KCl, 2 CaCl2, 8.2 MgCl2, 4 NaHCO3, 1 NaH2PO4-H20, 5 Trehalose, 10 Sucrose, 5 HEPES, pH=7.5. Brains were placed on a small glass coverslip (Carolina Cover Glasses, Circles, 12 mm, Cat. No: 633029) in a perfusion chamber filled with AHL. For P2X2 imaging, AHL buffer was perfused over the brains for 1 min of baseline GCaMP6 imaging, then ATP was delivered to the chamber by switching the perfusion flow from the channel containing AHL to the channel containing 2.5 mM ATP in AHL, pH 7.5. ATP was perfused for 2 min and then AHL was perfused for 2 min. Twelve-bit images were acquired with a 20 X inverted objective at 256 × 256-pixel resolution. Z-stacks were acquired every 5 sec for 3 min. Image processing and measurement of fluorescence intensity was performed in ImageJ as described previously^22^. For each cell body, fluorescence traces over time were normalized using this equation: ΔF/F = (F_n_-F_0_)/F_0_, where F_n_=fluorescence intensity recorded at time point n, and F_0_ is the average fluorescence value during the 1 min baseline recording. Maximum GCaMP change (ΔF/F) for individual cells was calculated using this equation: ΔF/F_max_ = (F_max_-F_0_)/F_0_, where F_max_=maximum fluorescence intensity value recorded during ATP application, and F_0_ is the average fluorescence value during the 1 min baseline recording. All analysis was done blind to experimental condition.

### Statistical analysis

All statistical analysis was done in GraphPad (Prism). For comparisons between 2 conditions, two-tailed unpaired *t*-tests were used. For comparisons between multiple groups, ordinary one-way ANOVAs followed by Tukey’s multiple comparison tests were used. For comparisons between different groups in the same analysis, ordinary one-way ANOVAs followed by Sidak’s multiple comparisons tests were used. For comparisons between time and genotype, two-way ANOVAs followed by Sidak’s multiple comparisons tests were used. \**P*<0.05, \*\**P*<0.01, \*\*\**P*<0.001. For all behavioral experiments, at least 3 technical replicates were performed. Representative confocal images are shown from at least 8-10 independent samples examined in each case. For live imaging, ∼15-20 neurons in 8-10 brains were used for each condition.

## Supplemental Figures and Figure Legends

**Figure S1:**
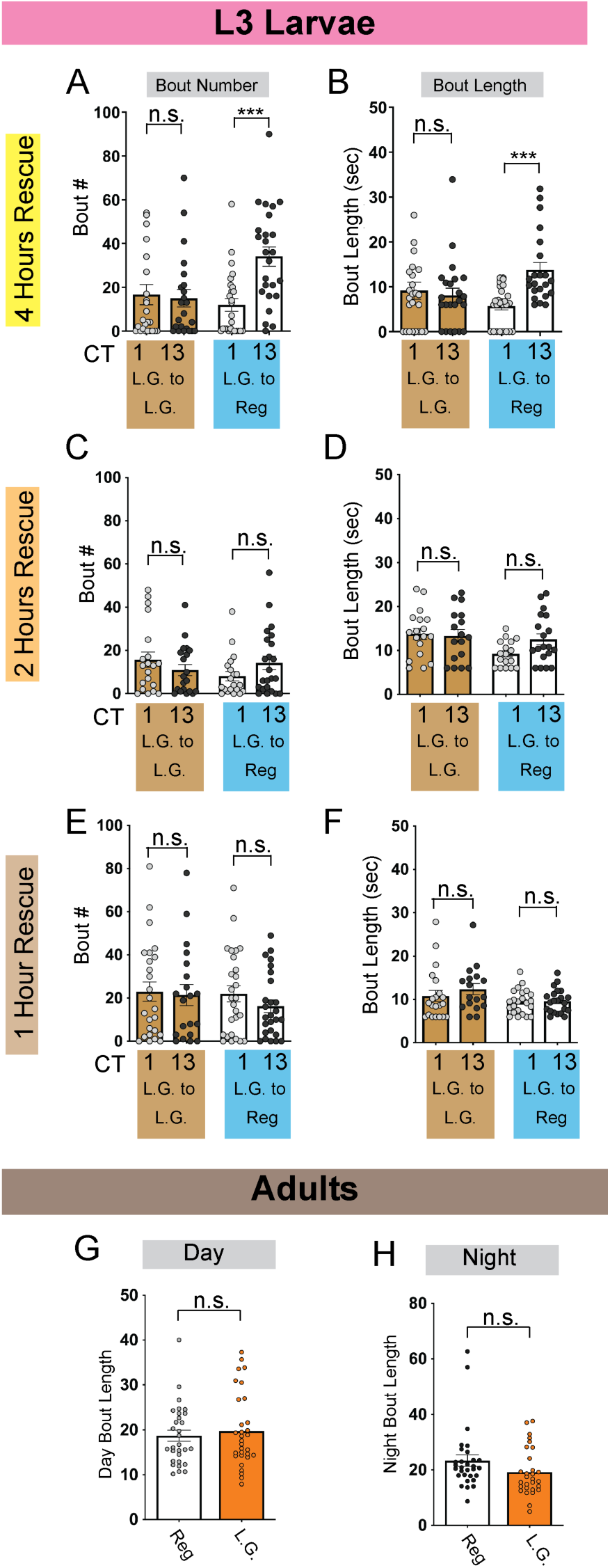
Sleep metrics in shifting nutrient conditions. (A-F) Sleep bout number (A, C, E) and bout length (B, D, F) at CT1 and CT13 in L3 raised on low glucose (L.G.) and moved to regular glucose diets (L.G. to Reg) or moved to L.G. (L.G. to L.G.) for 4 hrs (A and B), 2 hrs (C and D), and 1 hr (E and F). (G-L) Sleep measures for adult flies raised on regular glucose and placed on either low glucose (L.G.) or regular glucose (Reg) diets for 3 days. (G) Average daytime sleep bout duration. (H) Average nighttime sleep bout duration. A-F, n=18-25 larvae; G-H, n=28-30 flies. Two-way ANOVAs followed by Sidak’s multiple comparison test (A-F); unpaired two-tailed Student’s *t*-test (G-H).

**Figure S2:**
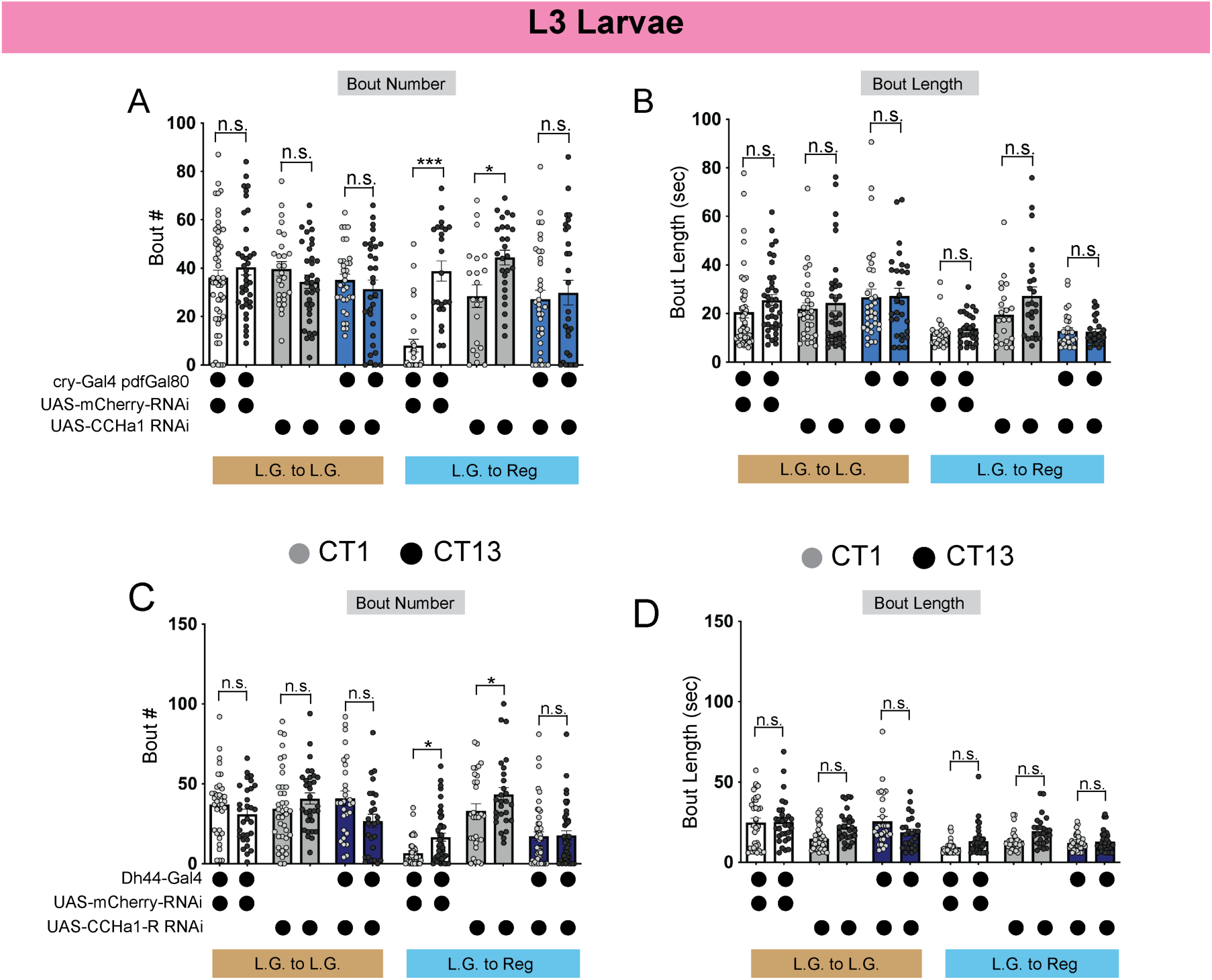
Sleep metrics for CCHamide-1 manipulations in L3 larvae. (A-B) Sleep bout number (A) and bout length (B) in L3 expressing UAS-CCHa1-RNAi with cry-Gal4 pdfGal80 and genetic controls at CT1 and CT13 in diet shifting paradigms. (C-D) Sleep bout number (C) and bout length (D) in L3 expressing UAS-CCHa1-R-RNAi with Dh44-Gal4 and genetic controls at CT1 and CT13 in diet shifting paradigms. Larvae were on either L.G. or Reg diets for 4 hours. A-B, n=25-47 larvae. Two-way ANOVAs followed by Sidak’s multiple comparison test (A-D).

**Figure S3:**
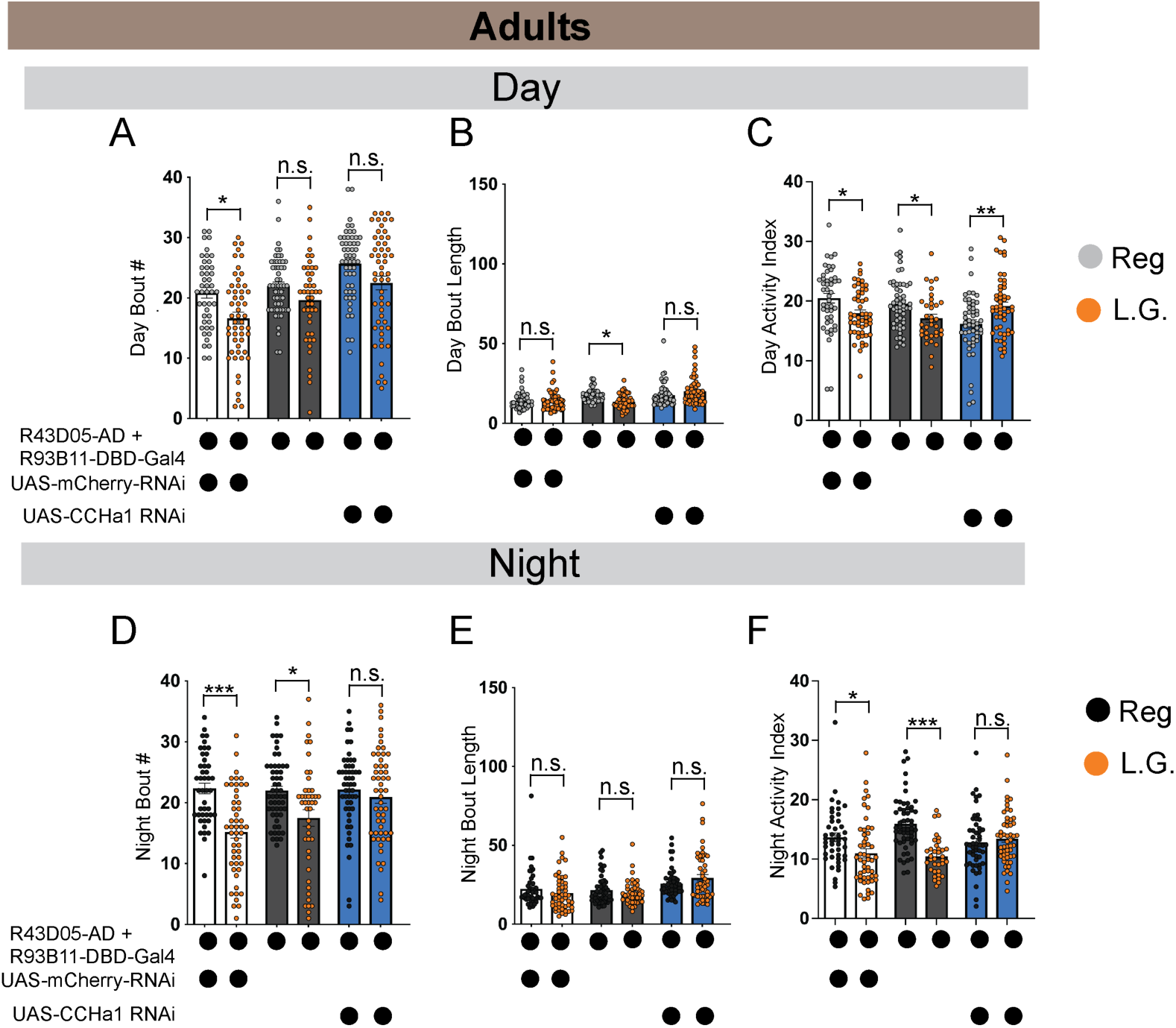
Sleep metrics for CCHamide-1 manipulations in adult flies. (A-F) Average number of daytime sleep bouts (A), average daytime sleep bout duration (B), daytime activity index (C), average number of nighttime sleep bouts (D), average nighttime sleep bout duration (E), and nighttime activity index (F) in adults expressing UAS-CCHa1-RNAi with DN1a-specific Gal4 and genetic controls. Orange dots indicate flies on L.G. diets. A-F, n= n=44-54 flies. One-way ANOVAs followed by Tukey’s multiple comparisons test (A-F).

**Figure S4:**
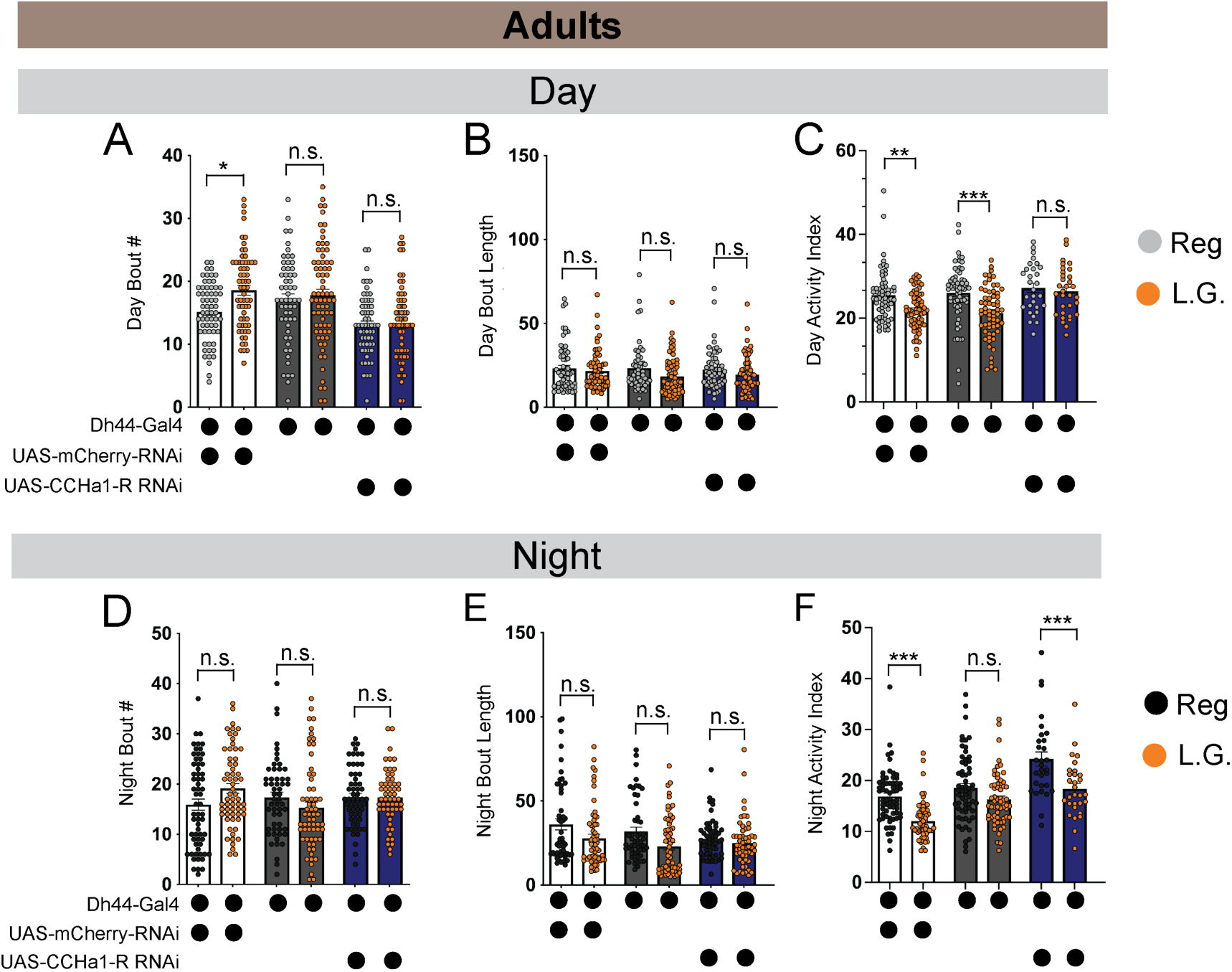
Sleep metrics for CCHamide-1 receptor manipulations in adult flies. (A-F) Average number of daytime sleep bouts (A), average daytime sleep bout duration (B), daytime activity index (C), average number of nighttime sleep bouts (D), average nighttime sleep bout duration (E), and nighttime activity index (F) in adults expressing UAS-CCHa1-R-RNAi with Dh44-Gal4 and genetic controls. Orange dots indicate flies on L.G. diets. A-F, n=55-64 flies. One-way ANOVAs followed by Tukey’s multiple comparisons test (A-F).

**Figure S5:**
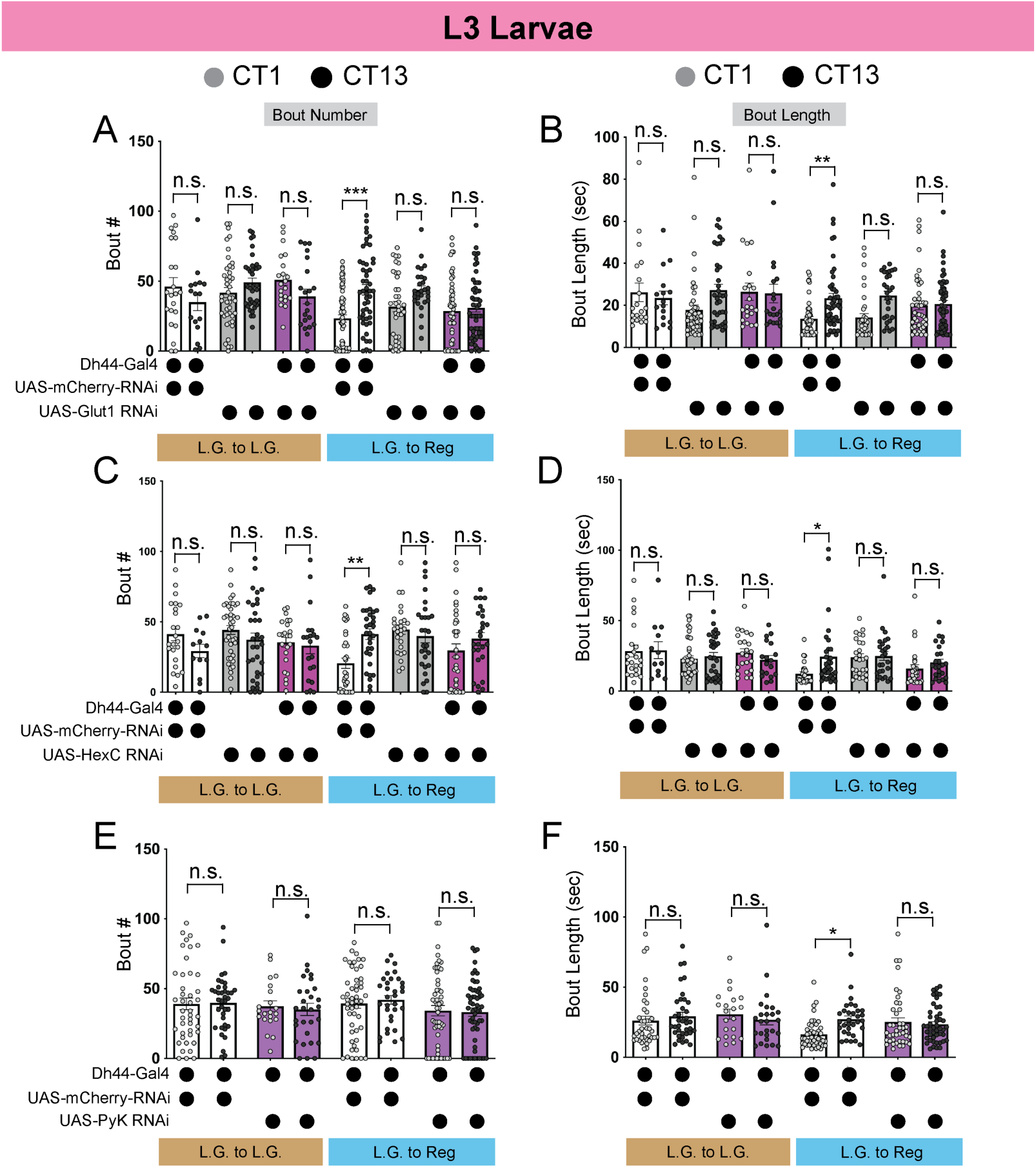
Sleep metrics for Dh44 glucose metabolism gene manipulations in L3 larvae. (A-B) Sleep bout number (A) and bout length (B) in L3 expressing UAS-Glut1-RNAi with Dh44-Gal4 and genetic controls at CT1 and CT13 in diet shifting paradigms. (C-D) Sleep bout number (C) and bout length (D) in L3 expressing UAS-HexC-RNAi with Dh44-Gal4 and genetic controls at CT1 and CT13 in diet shifting paradigms. (E-F) Sleep bout number (E) and bout length (F) in L3 expressing UAS-PyK-RNAi with Dh44-Gal4 and genetic controls at CT1 and CT13 in diet shifting paradigms. Larvae were on either L.G. or Reg diets for 4 hours. A-F, n=25-47 larvae. Two-way ANOVAs followed by Sidak’s multiple comparison test (A-F).

**Figure S6:**
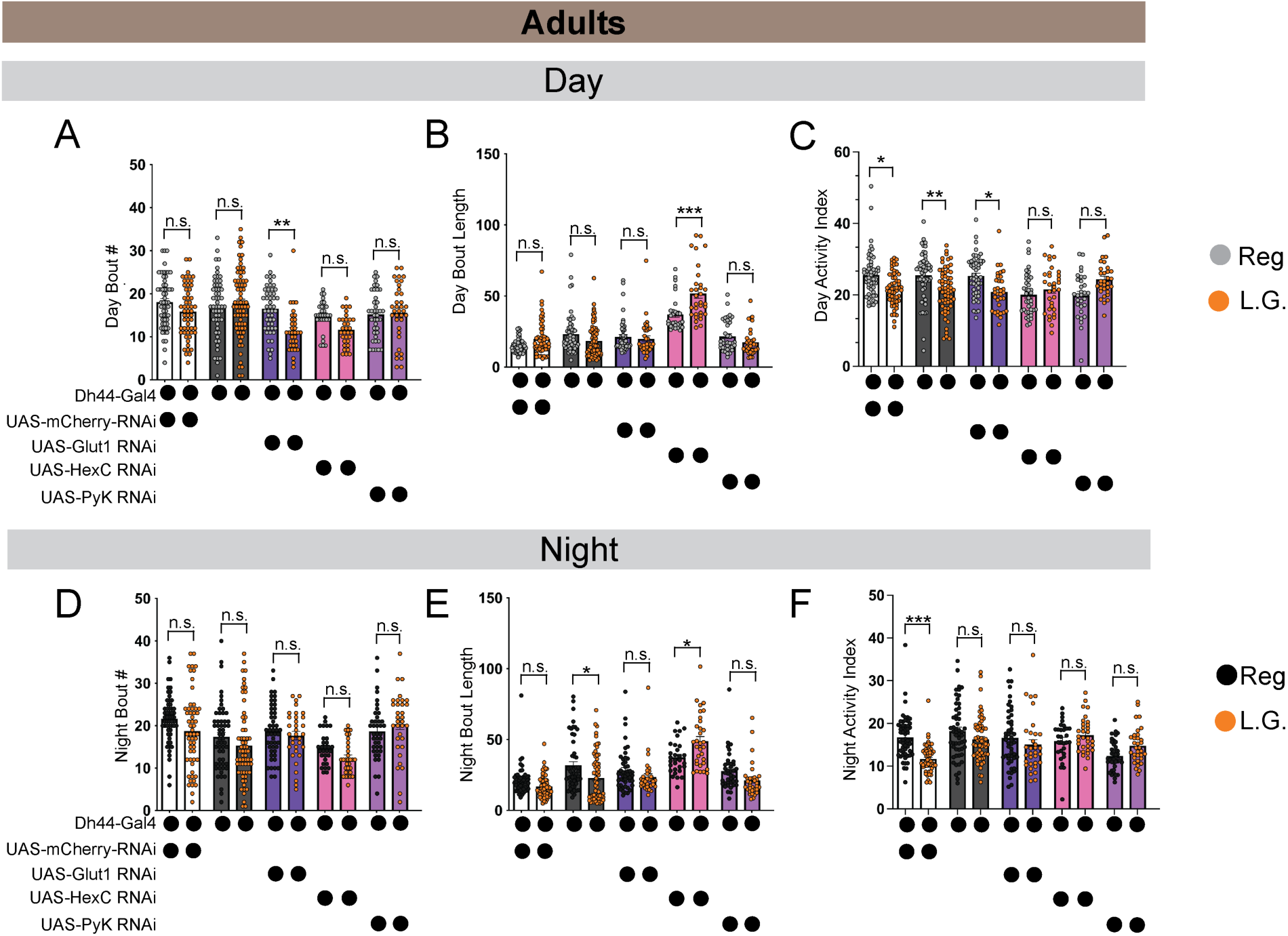
Sleep metrics for Dh44 glucose metabolism gene manipulations in adults. (A-D) Average number of daytime sleep bouts (A), average daytime sleep bout duration (B), and daytime activity index (C). Average number of nighttime sleep bouts (D), average nighttime sleep bout duration (E), and nighttime activity index (F) in adults expressing UAS-Glut1 RNAi, UAS-HexC RNAi, and UAS-PyK RNAi with Dh44-Gal4 and genetic controls. Orange dots indicate flies on L.G. diets. A-F, n=30-60 flies. One-way ANOVAs followed by Tukey’s multiple comparisons test (A-F).

**Figure S7:**
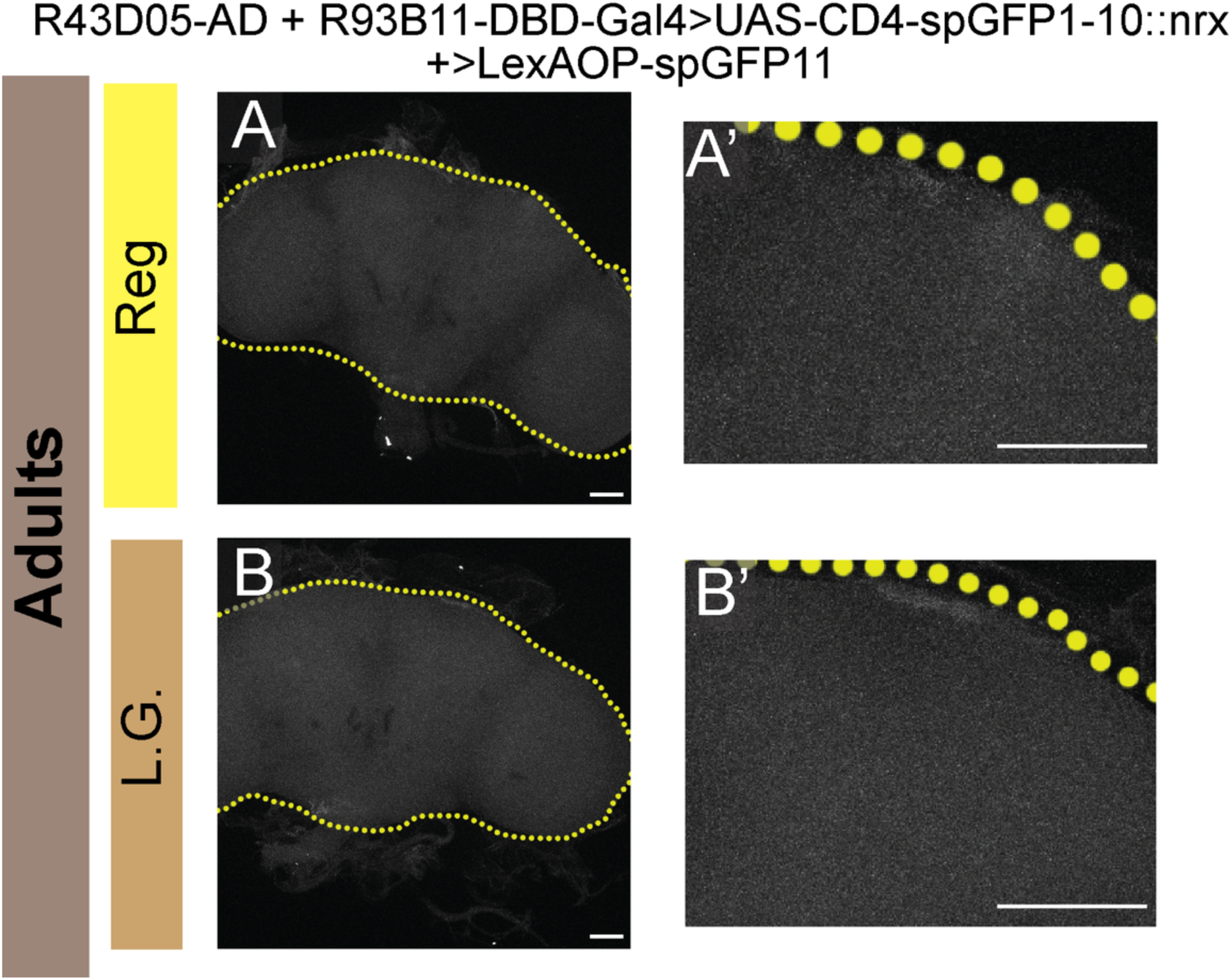
Confirmation of GRASP signal in adult brains. Neurexin-based GRASP with adult DN1a-Gal4 driver alone (R43D05-AD + R93B11-DBD) in flies on either regular glucose (A and A’) or L.G. (B and B’) diets. Higher magnification of region of interest in (A’) and (B’). Scale bars, 50 µm for A-B’.

